# Single-embryo and single-blastomere immunoblotting reports protein expression heterogeneity in early-stage preimplantation embryos

**DOI:** 10.1101/357731

**Authors:** Elisabet Rosàs-Canyelles, Andrew J. Modzelewski, Lin He, Amy E. Herr

**Author notes:** **Corresponding author**: Amy E. Herr.

## Abstract

Understanding how a zygote develops from a single cell into a multicellular organism has benefitted from single-cell tools, including RNA sequencing (RNA-Seq) and immunofluorescence (IF). However, scrutinizing inter- and intra-embryonic phenotypic variation is hindered by two fundamental limitations; the loose correlation between transcription and translation and the cross-reactivity of immunoreagents. To address these challenges, we describe a high-specificity microfluidic immunoblot optimized to quantify protein expression from all stages of mouse preimplantation development. Despite limited availability of isoform-specific immunoreagents, the immunoblot resolves inter-embryonic heterogeneity of embryo-specific isoforms (i.e., DICER-1). We observed significantly higher DICER-1 isoform expression in oocytes when compared to two-cell embryos, and further find that protein expression levels follow the same trend as mRNA for both the full-length and truncated DICER-1 isoforms. At the morula stage, we assayed both whole and disaggregated embryos for loading controls (β-tubulin, GAPDH) and markers that regulate cell fate decisions (CDX-2, SOX-2). In disaggregated morula, we found that cell volume showed positive, linear correlation with expression of β-tubulin and SOX-2. In dissociated two-cell and four-cell blastomeres, we detect significant inter-blastomeric variation in GADD45a expression, corroborating suspected cellular heterogeneity even in the earliest multicellular stage of preimplantation embryos. As RNA-Seq and other transcript-centric approaches continue to further probe preimplantation development, the demand for companion protein-based techniques rises. The reported microfluidic immunoblot serves as an essential tool for understanding mammalian development by providing high-specificity and direct measurements of protein targets at single-embryo and single-blastomere resolution.

## Introduction

The initiating events and specific proteins involved in the first cell fate commitment within pre-implantation blastomeres still remain open questions in developmental biology. While functional studies and embryonic plasticity suggest that blastomeres remain equivalent until the compacted morula^1–3^, growing evidence of inter-blastomeric differences in early-stage embryos point to heterogeneous configurations at even the earliest multicellular stages^4–14^. Although measurement tools with single-embryo and single-blastomere resolution, including RNA-Seq, have greatly advanced our knowledge, companion protein expression and state measurements within single embryos are required to test and validate these transcript-based predictions. Direct assessment of protein expression is required.

While immunofluorescence (IF) reports protein abundance and localization in embryos, IF is stymied by: (i) ubiquitous immunoreagent cross-reactivity that renders IF unsuitable for detection of small protein variations or multiplexing beyond ~5 targets^15^, (ii) proteoform ‘blind spots’ arising from limited isoform-specific immunoreagents that reduce the detectable repertoire of targets^16^, and (iii) confounding but required chemical fixation prior to IF measurement of endogenous intracellular proteins (i.e., epitope masking, cell morphology modifications, and perturbation of protein localization by diffusional gradients formed as fixation occurs)^17,18^. Flow cytometry and mass cytometry suffer from similar specificity and fixation concerns as IF19. Mass spectrometry does not yet afford the sensitivity to analyze single-mouse embryos or blastomeres (tens of ng of protein in oocytes to < 1 ng of protein in blastomeres at the blastocyst stage)^19,20^. Although protein analysis tools have been recently introduced to measure protein targets in single cells^21–24^, fundamental differences between cultured cell lines and mammalian embryos have hindered the study of early mammalian development. Key differences include cell size and composition, membrane structure and sample availability of ~20 embryos per mouse, equivalent to less than 1 μg of protein^25–28^. To complement the repertoire of existing measurements and resolve the intriguing questions surrounding mammalian development, such as when the first cell fate decisions are made, precision protein tools with higher selectivity are needed.

Here we report microfluidic immunoblotting for direct analysis of proteoforms across all stages of mouse preimplantation, from whole embryos to single blastomeres. With a dynamic detection range spanning two-orders of magnitude (oocytes at 10^−15^ mole to blastocyst blastomeres at 10^−17^ mole), we scrutinize widely used housekeeping proteins (β-tubulin, β-actinin, and GAPDH), embryo-specific isoforms (DICER-1) and regulators of cell fate decisions (SOX-2 and GADD45a). In whole morula, we assess expression of three targets (GAPDH, CDX-2 and SOX-2) having small 1–2 kDa differences in molecular mass. Applied to the study of intra-embryonic heterogeneity, we observe statistically significant differences in GADD45a expression at the four-cell and even two-cell stages. Microfluidic immunoblotting provides the resolution necessary to quantifiably investigate both inter- and intra-embryonic heterogeneity.

## Results

### Microfluidic immunoblotting of single embryos and single blastomeres

We first sought to directly measure protein expression in cells ranging from single oocytes (~80 μm in diameter) to single blastomeres from disaggregated blastocysts (<20 μm in diameter at 3.5–4.0 days post coitus, dpc) (Fig. 1a). Sample preparation of harvested murine embryos includes (i) removal of the *zona pellucida* by incubating with acidic Tyrode’s solution and, if studying disaggregated blastomeres, (ii) dissociation of embryos into individual blastomeres by incubation with trypsin and Accutase^®^. The microfluidic immunoblot, comprised of a 100–150 µm-thick polyacrylamide (PA) gel layered on a glass microscope slide, is stippled with an array of microwells (100–160 µm in diameter), where each microwell is designed to isolate an individual embryo or blastomere, depending on the assay. Using a standard mouth-controlled capillary tube assemblys^29^, single embryos or blastomeres are seated into individual microwells (Fig. 1b). Once isolated in each microwell, cell samples are chemically lysed (60–70 s). An electric field (E = 40 V/cm; 35–75 s) is applied to drive protein polyacrylamide gel electrophoresis (PAGE) in a 3 mm-long separation lane abutting each lysate-containing microwell. Protein blotting (immobilization) to the PA gel is achieved by UV-mediated activation of benzophenone methacrylamide moieties crosslinked into the gel matrix. Size-resolved immobilized protein bands are probed using primary antibodies and fluorophore-conjugated secondary antibodies, resulting in single-embryo or single-blastomere immunoblots. By probing for the protein loading control β-tubulin in lysate from single oocytes down to individual blastomeres from a disaggregated blastocyst (Fig. 1c), we determined a dynamic detection range spanning femtomoles (10^−15^) to tens of attomoles (10^−17^), assuming a starting protein target concentration in the μM range^30^.

**Fig. 1.**
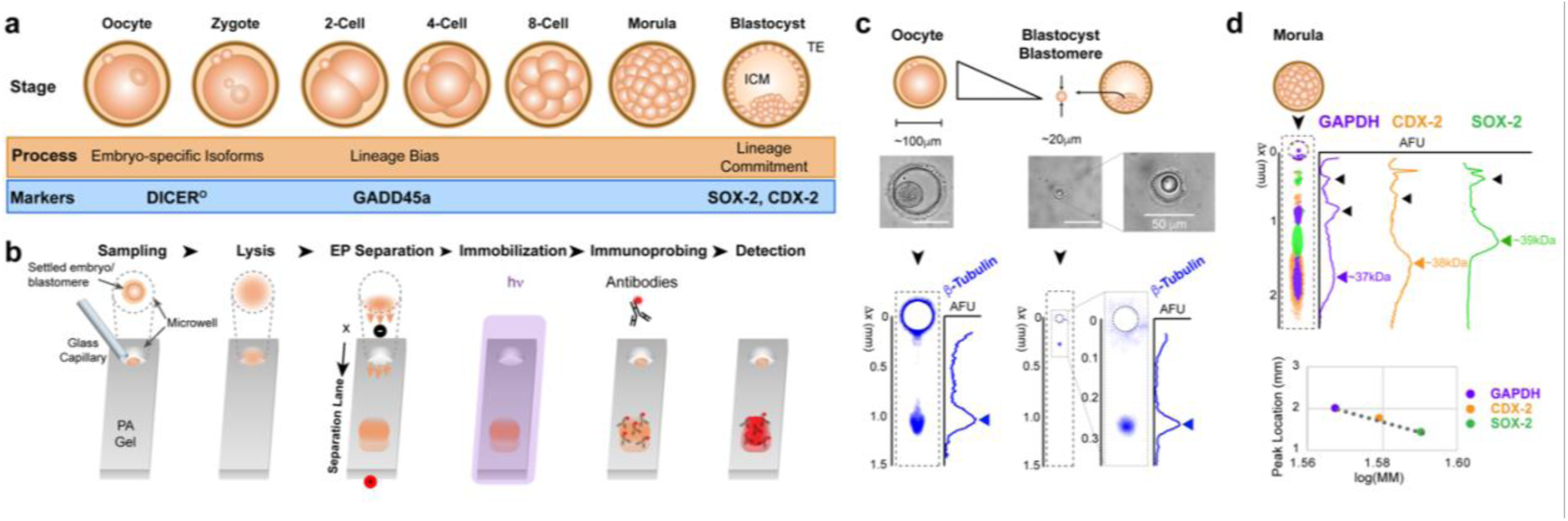
Microfluidic immunoblotting of single embryo and single blastomeres measures protein markers in all stages of preimplantation development. (a) During preimplantation development, a fertilized oocyte develops into a blastocyst. Protein markers investigated in this study are related to embryo-specific isoform expression, early-stage lineage biases and cell fate specification. (b) The microfluidic immunoblotting workflow begins with sampling a single embryo or single blastomere into a microwell patterned on a polyacrylamide (PA) gel. Samples are lysed and electrophoresed into the PA, achieving separation of proteins by molecular mass. Proteins are immobilized to gel matrix by UV-activated benzophenone chemistry and probed with fluorophore-conjugated antibodies. (c) Loading control β-tubulin was measured from single oocytes down to single blastocyst blastomeres. Brightfield micrographs of a settled oocyte and blastomere are shown above false-colored micrographs of resulting β-tubulin immunoblots and corresponding fluorescence intensity profiles. Arrows mark position of protein bands and scale bars are 100 μm, unless specified. (d) Single morula assayed for multiple targets that differ by 1–2 kDa (GAPDH, CDX-2 and SOX-2) show a strong log-linear relationship between migration distance and molecular mass (R^2^ = 0.9842).

We next scrutinized single morula (3.0 dpc) for SOX-2 and CDX-2, two transcription factors that regulate pluripotency and differentiation^31,32^, and the loading control GAPDH. SOX-2, CDX-2 and GAPDH have molecular masses of 37, 38 and 39 kDa, respectively (Fig. 1d). By employing a combination of (i) primary antibodies raised in different animals (goat-anti-GAPDH, rabbit-anti-SOX-2, and mouse-anti-CDX-2), (ii) secondary antibodies conjugated to different fluorophores (donkey-anti-goat, rabbit and mouse conjugated to AlexaFluor 555, 488 and 594, respectively), and (iii) reprobing after gel stripping using a reducing stripping buffer, the microfluidic immunoblot resolved all three targets from intact morula. The observed log-linear relationship between molecular mass and migration distance (Fig. 1d) enables distinguishing target protein bands from non-specific antibody signal and demonstrates that single-morula PAGE resolves protein targets with molecular mass as close as 1–2 kDa.

### Single-embryo and single-blastomere immunoblotting detects biological variation

Given our ability to immunoblot dissociated blastomeres, we next examined (i) if embryo disaggregation artificially alters the protein abundance of the whole embryo and (ii) if we can reconstruct the expression profile of the whole embryo, even when constituent blastomeres are assayed individually.

We first inquired if loading a pre-determined increase of protein in the microfluidic immunoblot would yield a concomitant increase in protein signal. We thus performed titrations where we controlled loaded protein by using individual blastomeres from dissociated four-cell embryos (2.0 dpc) as discrete and easily manipulable loads of protein. We loaded either one or two blastomeres into microwells and assayed the microwell lysate for β-tubulin (Fig. 2a). We observed a proportional increase in β-tubulin expression (area-under-the-curve signal or AUC) from microwells loaded with two blastomeres as compared to microwells loaded with one blastomere (Mann Whitney U Test, p value = 6.28×10^−5^, with *N* = 7 and 11 microwells, respectively). The observation corroborates the supposition that two blastomeres would contain two-fold more protein than a single blastomere (Fig. 2a).

**Fig. 2.**
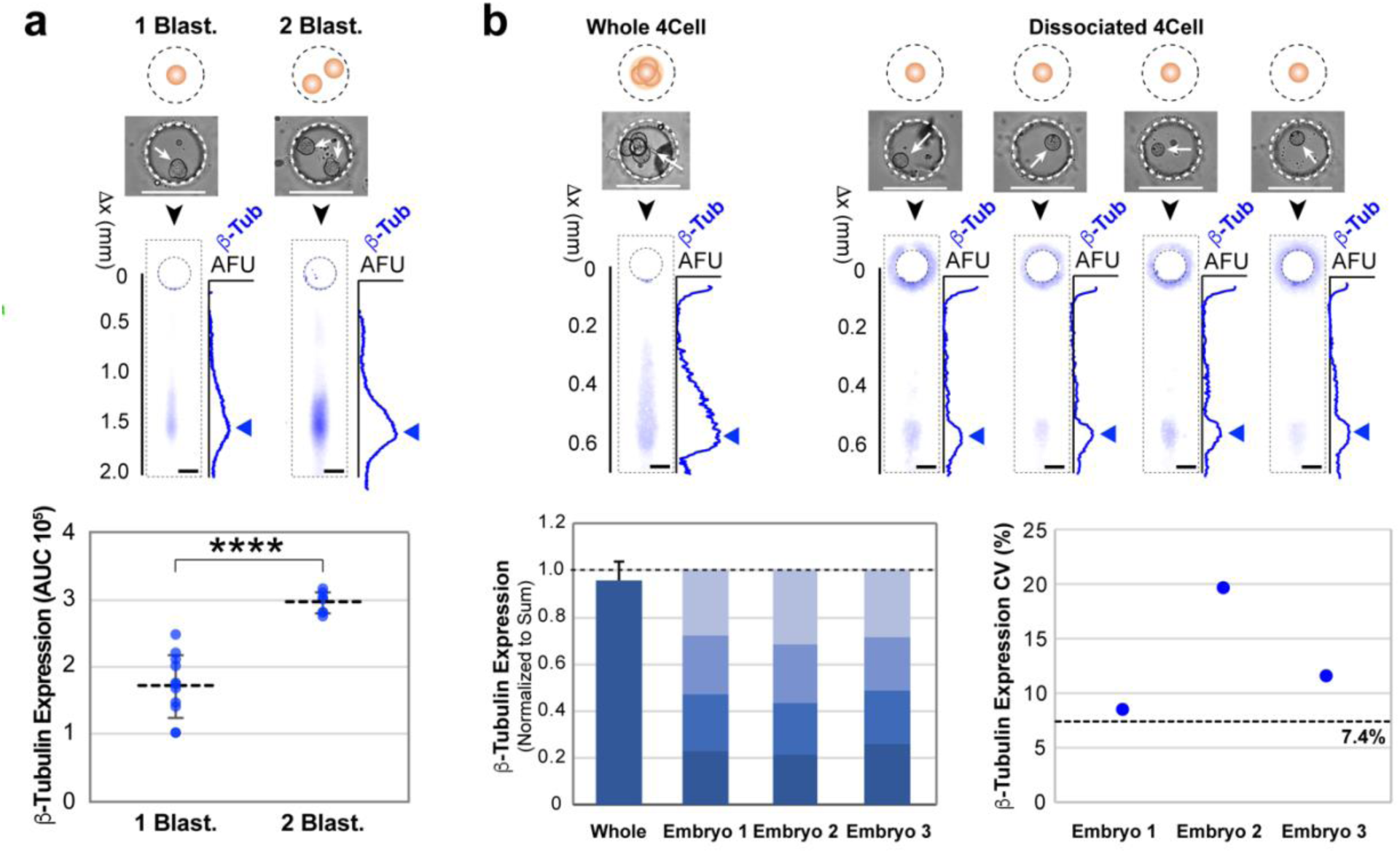
Microfluidic immunoblotting detects intra-embryonic biological variation in β-tubulin expression. (a) β-tubulin titration experiment. One or two blastomeres of dissociated four-cell embryos are sampled into microwells and assayed for β-tubulin. Brightfield images show blastomeres settled into microwells prior to lysis. Under these, false-color fluorescence micrographs and corresponding β-tubulin intensity profiles of resulting immunoblots. Arrows mark the position of protein bands. Dot plot of β-tubulin signal for immunoblots of one and two blastomeres demonstrate an increase in detection of β-tubulin for two blastomeres over one blastomere (horizontal bars indicate mean ± S.D., Mann Whitney U test, p value < 0.001, N = 11 and 7 for microwells with one and two blastomeres, respectively). (b) Reconstruction of whole embryo from disaggregated blastomeres. Bright field micrographs of whole and disaggregated four-cell embryos settled into microwells (top), with corresponding false-colored fluorescence micrographs of β-tubulin immunoblots. Intensity profiles are shown to the right of immunoblots, with blue arrow marking the position of the protein bands. Stacked bar graphs show individual blastomere contributions to total β-tubulin expression of four-cell embryos. Whole embryos assayed alongside dissociated blastomeres show similar levels of total β-tubulin expression (error bar indicates S.D., Wilcoxon matched-pairs signed rank test, p value = 0.5, *N* = 3 independent experiments), indicating sum of individually assayed blastomeres is equivalent to a whole embryo. Dot plot of β-tubulin expression coefficient of variation (CV %) for blastomeres of three disaggregated four-cell embryos. All CV values are above the technical CV_threshold_ of 7.4% (Fig. S1).

We next sought to assess if the protein profile of a whole embryo could be reconstructed from immunoblots of individual, dissociated blastomeres. To do so, we simultaneously assayed (i) intact individual four-cell embryos and (ii) blastomeres from disaggregated four-cell embryo (each blastomere contained in a separate microwell) (Fig. 2b). The resulting AUC for the β-tubulin protein band was compared between the intact four-cell embryo and the summed immunoblot signal from the four disaggregated blastomeres. We observed no significant difference between the sum of the contributions of dissociated blastomeres and signal obtained from a four-cell embryo (Wilcoxon matched-pairs signed rank test, p value = 0.5, *N* = 3 independent experiments) (Fig. 2b), suggesting that immunoblotting of individual blastomeres can reconstruct the protein profile of the originating intact four-cell embryo.

Finally, we sought to assess if the source of the observed inter-blastomeric variation in β-tubulin AUC was attributable to biological variation or confounding technical variation. First, we established a technical variation threshold by quantifying immunoblots of microwells uniformly loaded with purified protein. Briefly, we partitioned a solution of purified bovine serum albumin (BSA, 1 μM in PBS) into the microwells by incubating PA gels in BSA solution for 30 min. We then performed the immunoblotting assay and quantified BSA protein band AUC. We calculated the coefficient of variation in BSA AUC (CV_AUC_ % = AUC standard deviation (S.D.) / mean AUC x 100) and computed a technical variation threshold defined as > 3 x S.D. of the mean CV_AUC_^33^ (CV_threshold_ = mean CV_AUC_ + 3 S.D. = 7.4%, where mean CV_AUC_ = 4.69% and S.D. = 0.92%, Fig. S1). For all dissociated four-cell embryos studied, the inter-blastomeric β-tubulin expression CV exceeded the technical variation threshold (CVs = 8.3%, 19.6% and 11.3% for embryos, Fig. 2b). Consequently, we attribute the inter-blastomeric variation to biological variation and not technical variation.

### Normalization of SOX-2 expression by β-tubulin is not equivalent to normalization by cell volume for morula blastomeres

RNA-Seq studies suggest that cells regulate transcription to maintain mRNA concentration in response to changes in cell volumes^34,35^. Thus, cellular mRNA concentration is a more accurate indicator of cell phenotype than cellular mRNA abundance^34,35^. Normalizing by a loading control that is strongly correlated with cell size is therefore crucial for elucidating true phenotypic differences between cells. However, RNA-based studies show that commonly employed housekeeping genes (e.g., GAPDH and β-tubulin) are not stably and homogeneously expressed across different samples, experimental conditions or treatments^36^. The issue is further exaggerated as the sample size diminishes, reducing the averaging effect of a larger pooled samples. Whether this variability, and thus unreliability, of loading controls for single-cell studies prevails at the protein level remains to be explored.

Hence, we next tested whether the widely-used loading control β-tubulin is an accurate indicator of cell volume in preimplantation embryos. We assayed dissociated morula blastomeres for β-tubulin and compared β-tubulin expression (AUC) to cell volume (computed from brightfield images of cells seated in microwells) (Fig. 3a). We noted a significant positive, linear correlation between cell volume and β-tubulin expression (Pearson correlation, ρ = 0.8582, N = 8, p value = 0.0064) (Fig. 3b). To determine if this correlation indicates that β-tubulin can be used as a proxy for cell volume, we examined whether normalizing SOX-2 expression by β-tubulin expression is equivalent to normalizing by computed cell volume. We first observed that while SOX-2 expression showed significant positive correlation with cell volume (Pearson correlation, ρ = 0.7660, N = 8, p value = 0.0267) (Fig. 3b), SOX-2 expression normalized by β-tubulin was not correlated with cell volume (negative association, Pearson correlation ρ = −0.5866, N = 7, p value = 0.1668) (Fig. 3b). Furthermore, SOX-2 normalized by cell volume was not significantly correlated with SOX-2 normalized by β-tubulin (Pearson correlation, ρ = 0.5522, p value = 0.1558) (Fig. 3b).

**Fig. 3.**
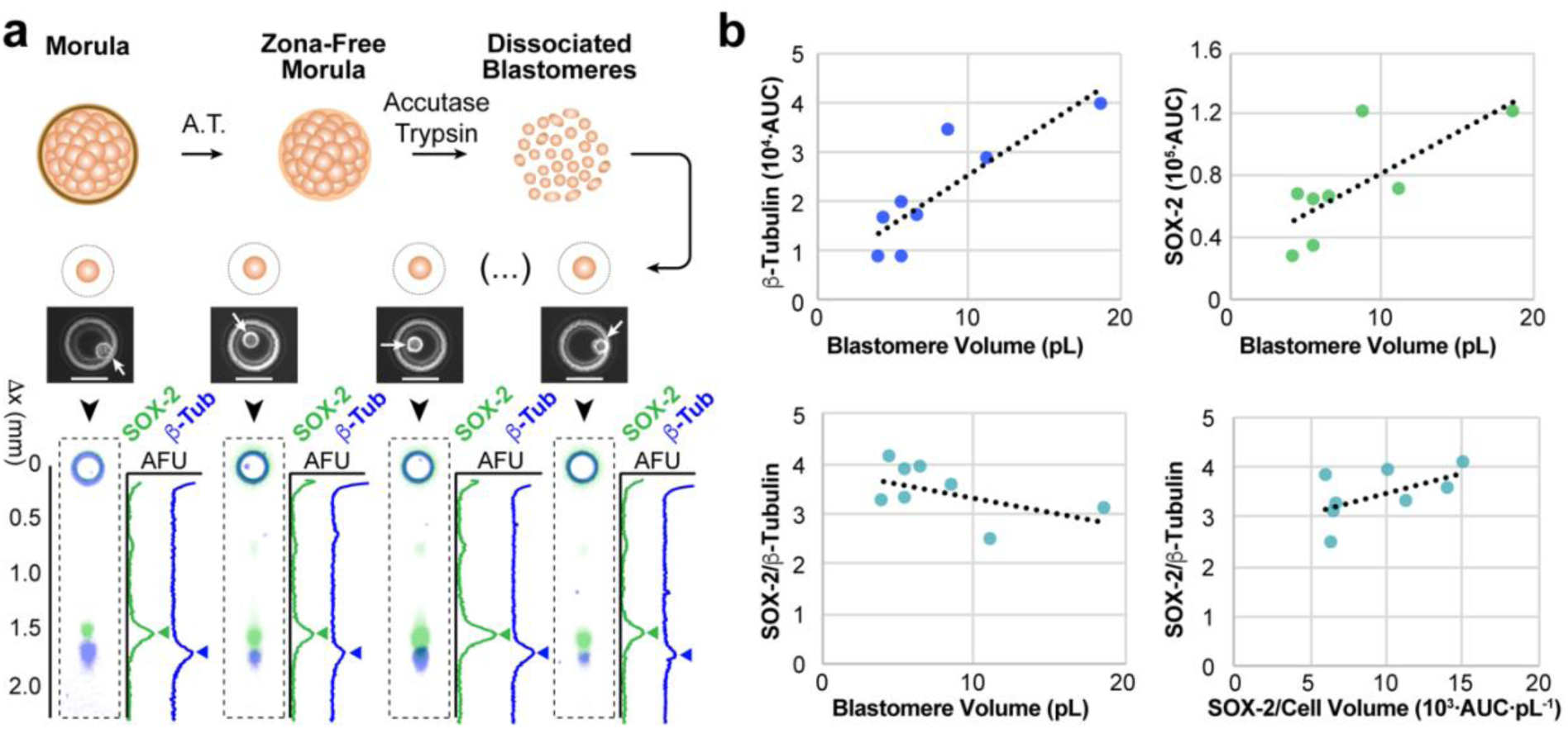
Single-blastomere immunoblotting identifies correlations between cell volume and marker expression in dissociated morula blastomeres. (a) Immunoblotting dissociated morula blastomeres for β-tubulin and SOX-2. Schematic (top) for dissociation of whole morula into individual blastomeres, which are seated into microwells of an immunoblotting device as shown in bright field images. False-colored fluorescence micrographs show β-tubulin and SOX-2 protein bands, with intensity profiles shown adjacent to micrographs. Arrows mark the position of protein bands. (b) Bivariate plot of blastomere cell volume and β-tubulin expression shows significant positive linear correlation (Pearson correlation, N = 8, ρ = 0.582, p value = 0.0064). While cell volume also shows significant positive linear correlation with SOX-2 expression, the same is not true when normalizing SOX-2 expression by β-tubulin expression (Pearson correlation, N = 8, ρ = 0.7381 and −0.5232 with p values = 0.0366 and 0.1835, respectively). Bivariate plot of SOX-2 expression normalized by β-tubulin and by cell volume shows a positive, but non-significant, association (Pearson correlation, N = 8, ρ = 0.5522, p value = 0.1559).

Hence, careful validation of loading controls as accurate predictors of cell volume is indispensable, even if expression of loading controls is well correlated with cell volume. Indexing endpoint immunoblot results with micrographs of the originating and intact cell sample allows us to normalize target protein expression by cell volume – a normalization that is impossible in endpoint immunoblots of lysate from pooled cell populations.

### Microfluidic immunoblotting detects truncated DICER-1 isoform expression in oocytes and two-cell embryos

Alternative splicing is frequent during early embryonic development in mouse and human^37–39^. However, efforts to investigate whether the corresponding alternate protein isoforms are ultimately and stably generated require pooling tens of thousands of collected embryos from each stage, losing intra-blastomeric information in the process^20^. Thus, single embryo and blastomere approaches capable of resolving proteoforms resulting from alternative splicing are needed.

To this end, we aimed to examine one of the earliest known examples of a protein isoform that exists in mouse development. DICER-1 is essential for small RNA-mediated gene expression regulation. By processing small RNAs into their mature form, DICER-1 generates the sequence-specific guides required by effector complexes to target cognate mRNAs and repress their translation ^40^. Bulk analyses of mouse oocytes found high expression of an N-terminally truncated isoform, denoted DICERO (Fig. 4a). DICERO demonstrates higher catalytic activity than its full-length form (DICER-1) and is believed to drive the high activity of endogenous small interfering RNAs (endo-siRNAs) in mouse oocytes, but not in somatic cells^40^. The DicerO transcript persists until the fertilized zygote stage, but it remains unclear whether the DICERO protein isoform is exclusive to oogenesis or is maternally inherited by the preimplantation embryo.

**Fig. 4.**
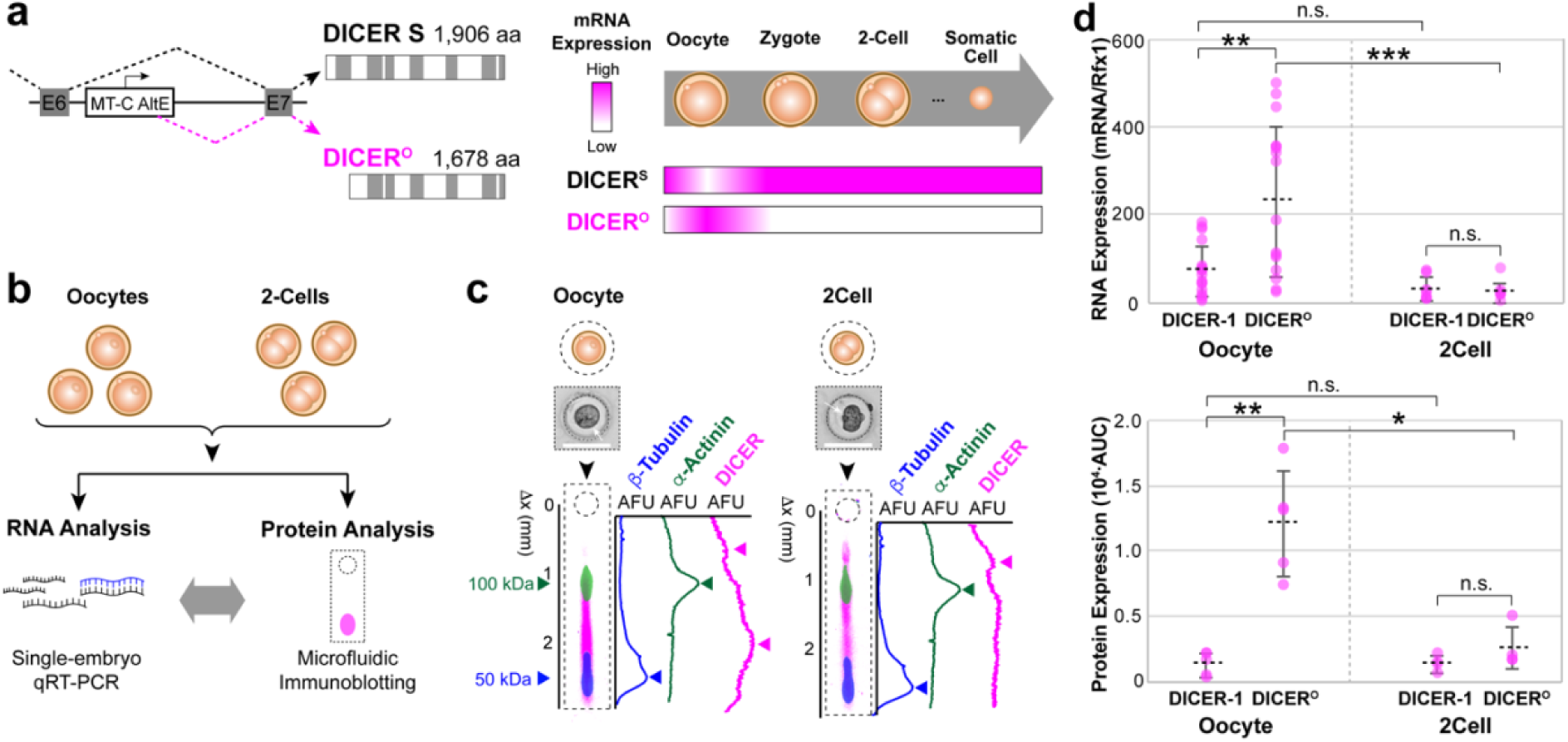
Higher DICER-1 isoform expression in oocytes than in two-cell embryos correlates with mRNA levels. (a) DICERO, a truncated isoform of DICER-1, appears only at the oocyte stage and is a product of alternative promoter usage. (b) Schematic of oocytes and two-cell embryos analyzed either by microfluidic immunoblotting or by companion qRT-PCR analysis. (c) Bright field micrographs of a settled oocyte and two-cell embryo. Under these, corresponding overlaid false-colored fluorescence micrographs and intensity profiles show protein bands for loading controls (β-actinin and β-tubulin) and DICER-1, where oocyte immunoblot demonstrates presence of a full-length DICER-1 (top arrow) and a lower molecular weight isoform (bottom arrow). Scale bars are 100 μm. (d) Dot plots of DICER isoform mRNA levels normalized by endogenous control Rfx1 (top) and protein expression (AUC, bottom) for single oocytes and single two-cell embryos. Expression of the truncated isoform is higher than the full-length DICER-1 for both mRNA andprotein in oocytes (mRNA/Rfx1_DICER-1_ vs. mRNA/Rfx1_DICER_°: Mann-Whitney U test, p value = 0.0052for N = 18; for AUC_DICER-1_ vs. AUC_DICER_°: Mann-Whitney U test, p value = 0.0079 for N = 5), but notin two-cell embryos (for mRNA/Rfx1_DICER-1_ vs. mRNA/Rfx1_DICER_°: Mann-Whitney U test, p value = 0.9551 for N = 7 for DICER-1 and N = 8 for DICER°; for AUC_DICER-1_ vs. AUC_DICER_°: Mann-Whitney U test, p value = 0.20 for N = 4). Oocytes show higher mRNA and protein expression than two-cells for the truncated isoform (mRNA/Rfx1_DICER_°: Mann Whitney U test, p value = 0.0004 for N = 18 oocytes and 8 two-cells; AUC_DICER_°: Mann Whitney U test, p value = 0.0159 for N = 5 oocytes and 4 two-cell embryos), but not the full-length DICER-1 (mRNA/Rfx1_DICER-1_: Mann Whitney U test, p value = 0.084 for N = 18 oocytes and 7 two-cell embryos; AUC_DICER-1_: Mann Whitney U test, p value= 0.9048 for N= 5 oocytes and 4 two-cell embryos) (horizontal bars indicate mean ± S.D.).

To explore whether DICERO is specific to the oocyte stage, we assayed oocytes and two-cell embryos for isoforms of DICER-1. We collected oocytes and two-cell embryos and divided each sample for analysis of either protein by microfluidic immunoblotting or mRNA analysis by single-embryo quantitative reverse transcription polymerase chain reaction (qRT-PCR) (Fig. 4b). Despite of a lack of an isoform-specific antibody, the electrophoretic separation of proteins resolved multiple DICER-1 isoforms by molecular mass. We observed that both oocytes and two-cell embryos expressed both the full-length and the truncated DICER-1 (Fig. 4c). For oocytes, we observed significantly higher expression of the truncated isoform over the full-length DICER-1 for both mRNA (normalized by endogenous control Rfx1) and protein (AUC) (for mRNA/Rfx1_DICER-1_ vs. mRNA/Rfx1DICER°: Mann-Whitney U test, p value = 0.0052 for N = 18; for AUCDICER-1 vs. AUCDICER°: Mann-Whitney U test, p value = 0.0079 for N = 5) (Fig. 4d). On the other hand, we found no significant difference in expression of truncated and full-length isoforms of DICER-1 in two-cell embryos (for mRNA/Rfx1DICER-1 vs. mRNA/Rfx1DICER°: Mann-Whitney U test, p value = 0.9551 for N = 7 for DICER-1 and N = 8 for DICER^o^; for AUCDICER-1 vs. AUCDICER°: Mann-Whitney U test, p value = 0.20 for N = 4) (Fig. 4d). When comparing expression levels between embryonic stages, we observed that for both mRNA and protein the expression of full-length DICER-1 was not significantly different between oocytes and two-cell embryos (mRNA/Rfx1DICER-1: Mann Whitney U test, p value = 0.084 for N = 18 oocytes and 7 two-cell embryos; AUCDICER-1: Mann Whitney U test, p value = 0.9048 for N = 5 oocytes and 4 two-cell embryos) (Fig. 4d). For the truncated isoform, however, we observed a significant decrease in both mRNA levels and protein levels from the oocyte to the two-cell stage (mRNA/Rfx1_DICER_°: Mann Whitney U test, p value = 0.0004 for N = 18 oocytes and 8 two-cells; AUC_DICER_°: Mann Whitney U test, p value = 0.0159 for N = 5 oocytes and 4 two-cell embryos) (Fig. 4d). Hence, protein PAGE from single-embryo lysates grants the selectivity required for measuring protein isoforms, even when pan-specific antibodies are the only reagent available.

### Single-blastomere immunoblotting reports GADD45a expression heterogeneity in two- and four-cell embryos

We finally sought to inspect early-stage embryos for lineage biases by measuring protein expression from disaggregated two-cell and four-cell embryos. The exact stage and circumstances by which blastomeres acquire certain fates remains unknown. On the one hand, it is thought that embryonic plasticity supports blastomere symmetry up to the 8-cell embryo, where embryos can compensate for the loss of one blastomere as early as the two-cell stage^41^. On the other hand, studies point to early asymmetry, where sister blastomeres show consistent bimodal expression of genes related to differentiation, suggesting that the involved factors may not be inherited equally by all blastomeres^6^.

As such, to quantitatively examine intra-embryonic heterogeneity in cell fate related markers, we assayed early-stage blastomeres for GADD45a, a protein involved in DNA damage repair that has been reported to show bimodal transcription at the two-cell and four-cell stages4 (Fig. 5a). We compared the intra-embryonic heterogeneity of GADD45a expression to that of β-tubulin, to control for stochasticity of protein partitioning at cell division ^42^ (Fig. 5b). We observed that the intra-embryonic variation in GADD45a expression is significantly higher than the variation in β-tubulin expression (CV_GADD45a_ = 31.5 ± 13.5%; CV_b-tub_ = 13.4 ± 5.4%; mean ± S.D., Wilcoxon signed-rank test, p value = 0.0312, *N* = 6, where all CVs > CV_threshold_ of 7.4%, indicating that the measured heterogeneity is attributable to biological, not technical, variation) (Fig. 5c & d). These findings suggest that blastomeres of four-cell embryos show heterogeneous expression of GADD45a, in agreement with the mRNA and IF-based findings of Biase *et al.*^4^.

**Fig. 5.**
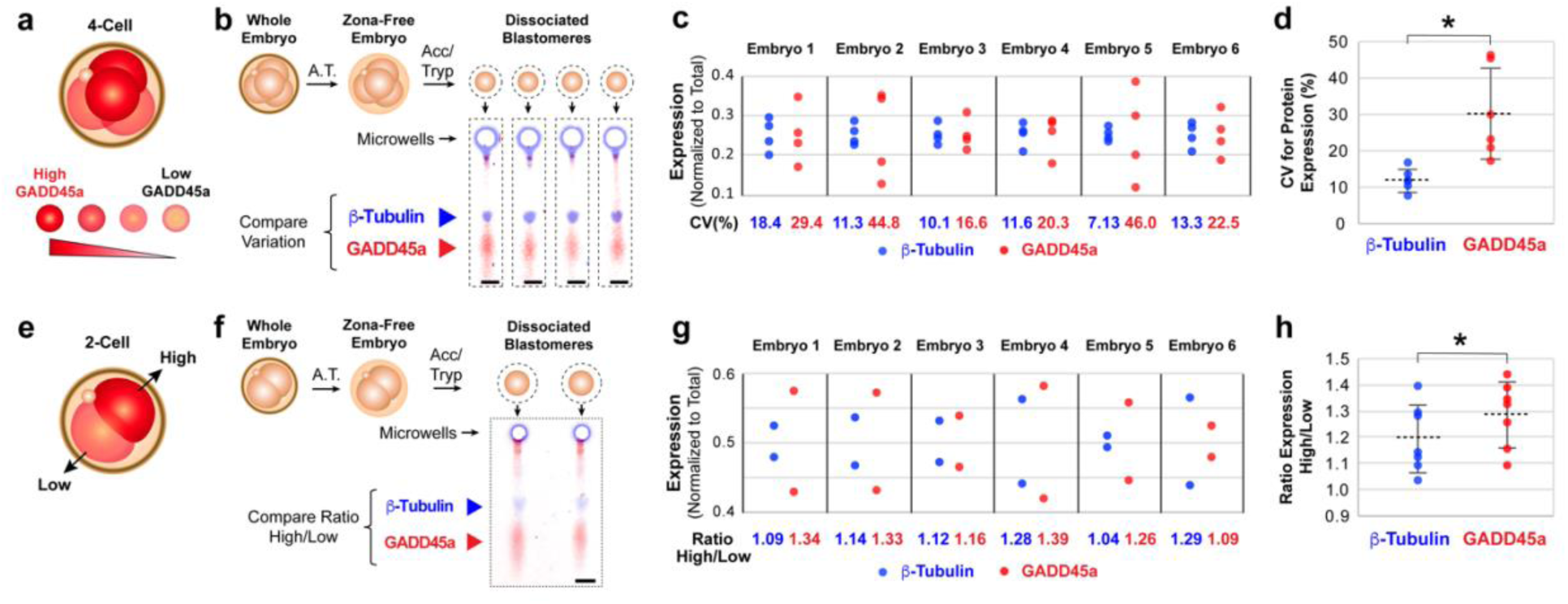
Microfluidic immunoblotting measures intra-embryonic heterogeneity in GADD45a expression in four-cell and two-cell embryos. (a) Four-cell embryos have been suggested to show early lineage bias by heterogeneous expression of GADD45a. (b) Design for testing this hypothesis includes removing *zona pellucida* from four-cell embryos and dissociation into individual blastomeres for subsequent immunoblotting for GADD45a and loading control β-tubulin. False-colored fluorescence micrographs show bands for both protein targets for one dissociated four-cell embryo. (c) Dot plot of expression of β-tubulin (blue) and GADD45a (red) by individual blastomeres, normalized to total expression, with corresponding inter-blastomeric coefficient of variation (CV) for β-tubulin and GADD45a expression. (d) Dot plot of CVs shows that inter-blastomeric variation in GADD45a is significantly different from inter-blastomeric variation in β-tubulin (Wilcoxon signed-rank test, p value = 0.0312, *N* = 6). We investigated whether this heterogeneity can be traced back to the two-cell stage (e). (f) Schematic of two-cell embryo sample preparation by removal of *zona pellucida* and dissociation into individual blastomeres. False-colored fluorescence micrographs show β-tubulin and GADD45a immunoblots for two-cell sister blastomeres. (g) Dot plots of β-tubulin and GADD45a expression by sister blastomeres, normalized to sum of expression of sister blastomeres. (h) Dot plot of ratio between the AUC of high-expressing and low-expressing blastomeres for β-tubulin and GADD45a (paired t test p value= 0.0251, N = 11 dissociated two-cell embryos) (horizontal bars in (d) and (h) indicate mean ± S.D.).

We next investigated if heterogeneity in GADD45a expression arises in the earlier two-cell embryo. Unlike in the four-cell stage, bimodality in GADD45a protein expression at the two-cell stage remains unexplored. We thus assayed dissociated two-cell embryos to understand the intra-embryonic distribution of GADD45a. To test whether one blastomere consistently showed higher GADD45a AUC than the other, we immunoblotted dissociated two-cell blastomeres for GADD45a and β-tubulin (Fig. 5e and f) and computed the ratio of expression between the blastomere with high expression and low expression for both markers (Fig. 5g). We found that the ratio of high-to-low GADD45a expression (1.26 ± 0.109) is significantly larger than the high-to-low ratio for β-tubulin (1.16 ± 0.11; mean ± S.D., paired t test p value = 0.0251, N = 11 two-cell embryos, distributions for GADD45a and β-tubulin passed the D’Agostino & Pearson normality test with p values = 0.6842 and 0.2497, respectively) (Fig. 5h). Hence, we posit that two-cell blastomeres display significant heterogeneity in GADD45a protein expression, providing protein-based evidence for heterogeneity in GADD45a expression even at the earliest multicellular preimplantation stage. Functional studies will allow us to determine whether differences in GADD5a protein expression levels are indicative of differential cellular phenotypes.

## Discussion

Open questions surround the timing and mechanism by which the first cell fate decisions are made during mammalian pre-implantation development. Do developing blastomeres remain homogeneous and functionally equivalent until the compacted morula stage ^1,2^? Or, alternately, do these developing blastomeres exhibit symmetry-breaking heterogeneous configurations, perhaps as early as the two-cell stage ^7,9–11^? In mice, zygotic genome activation (ZGA) occurs shortly after fertilization and is not fully realized until the two-cell stage, at which point nascent mRNA populate the embryonic transcriptome ^43^. While recent single-cell RNA-Seq experiments reveal sister blastomere transcriptome heterogeneity as early as the two-cell embryo in both human and mice^4,11^, functional studies suggest these differences may not matter until the four-cell stage^3,12^.

Here, microfluidic design provides an avenue for a cellular-resolution form of protein immunoblotting applicable to mammalian development as early as the oocyte stage of a murine model. To advance linking mRNA levels to protein expression when tackling questions on cell fate determination, we scrutinized two-cell and four-cell embryos for signs of heterogeneity in protein expression of GADD45a, a gene shown to be bimodally transcribed in early embryonic stages. The microfluidic immunoblot detects higher heterogeneity in GADD45a expression than loading control β-tubulin between the blastomeres in early embryos, providing the first protein-based validation of recent single cell RNA-Seq predictions. Companion functional competency measurements guided by the ever-increasing resolution of single-cell transcript approaches will help in determining the proteins and isoforms involved in key cell fate decisions event.

Directly scrutinizing and independently validating single-cell transcriptional data describing early embryonic development requires direct measurement of proteins with single-cell resolution. The asynchrony observed between mRNA and protein expression may simply reflect the uncoupled relationship between transcription and translation in the early embryo (~15 hr), synchronizing more tightly in later cleavage stages ^44^. Transcript abundance is not an accurate determinant of protein abundance^45,46^, and whether the required activating post translational modifications (PTMs) are present has been difficult to ascertain^47^. Beyond PTMs and protein-mediated signaling, isoforms of proteins such as DICER1 are involved in regulation of small RNA-mediated gene expression. Microfluidic immunoblotting resolved DICER-1 isoforms in single oocytes and two-cell embryos, with the truncated DICER-1 protein isoform as the dominant isoform in the oocyte stage. Significantly lower abundance of the truncated DICER-1 mRNA and protein in two-cell embryos compared to oocytes suggests that these may be inherited from the oocyte stage. These findings support previous studies suggesting that the truncated DICER-1 isoform, which shows higher activity than the full-length DICER-1, is oocyte-specific. Previously unattainable single-embryo lysate protein separations enable resolving and quantifying protein isoforms, even when isoform-specific antibodies are not available.

As detailed here, the ~20 embryos harvested from a single mouse donor are sufficient not just for one immunoblot, but for multiple single-embryo and single-blastomere immunoblots. The precision in sample handling and in enhanced sensitivity notably reduces the conventional PAGE sample requirements of several hundreds or thousands of embryos^20,40^. The implications are multi-fold. First, as single-embryo immunoblots inherently and dramatically lower sample requirements, the burden of animal sacrifice is likewise reduced. Second, statistical interpretation of single-embryo and single-blastomere immunoblot results is feasible, revealing intra-embryonic heterogeneity, as well as significant differences between embryos of the same fertilization event and between donors.

Lastly arises the possibility of companion immunoblot and mRNA assays on embryos of the same donor, thus enhancing the biological accuracy of correlations between mRNA levels and protein expression at different stages of the preimplantation embryo. Such insight into the expression dynamics would clarify how modulation in transcription dictates cellular phenotype ^48^. Moreover, with the advent of new gene editing technologies, (e.g., CRISPR, genomic screening methods including targeted, exome or whole genome sequencing) screening for off-target activity has become critical ^49^. Protein assays that can complement genomic screening, such as the one described in this study, will be crucial for screening embryos for protein-level effects of both on-target and off-target mutations, even when the latter occur in non-coding regions.

## Methods

### Animals and Ethics Statement

As a matter of caution and compliance, all appropriate authorizations have been acquired from institutional and/or federal regulatory bodies prior to performing this protocol. All mouse use, including but not limited to housing, breeding, production, sample collection for genotyping, and euthanasia, is in accordance with the Animal Welfare Act, the AVMA Guidelines on Euthanasia and are in compliance with the ILAR *Guide for Care and Use of Laboratory Animals,* and the UC Berkeley Institutional Animal Care and Use Committee (IACUC) guidelines and policies. Our animal care and use protocol has been reviewed and approved by our IACUC for this project.

### Chemical Reagents

Tetramethylethylenediamine (TEMED, T9281), ammonium persulfate (APS, A3678), β-mercaptoethanol (M3148), 30%T/2.7%C acrylamide/bis-acrylamide (37.5:1) (A3699), bovine serum albumin (BSA, A9418), Tyrode’s solution (T1788), trypsin 10X (59427 C), Accutase^®^ (A6964) and 3-(trimethoxylsilyl)propyl methacrylate (440159) were purchased from Sigma-Aldrich. Triton X-100 (BP-151) and phosphate-buffered saline (PBS, pH 7.4, 10010023) were purchased from Thermo Fisher Scientific. Premixed 10X tris-glycine EP buffer (25 mM Tris, pH 8.3; 192 mM glycine; 0.1% SDS) was purchased from Bio-Rad. Tris buffered saline with Tween-20 (TBST) was prepared from 20X TBST (sc-24953, Santa Cruz Biotechnology, Dallas, TX). Deionized water (18.2 MΩ) was obtained using an Ultrapure water system from Millipore. Alexa555-labeled bovine serum albumin (A34786) was purchased from Invitrogen. N-[3-[(3-Benzoylphenyl)formamido]propyl] methacrylamide (BPMAC) was custom synthesized by Pharm-Agra Laboratories (Brevard, NC). Gel Slick^TM^ (50640) was purchased from Lonza.

### Device Fabrication

Devices were fabricated using SU-8 wafers as previously reported. Microwell height and diameter were varied to accommodate different embryo stages (microwell diameters ranged from 20 μm for dissociated blastocyst blastomeres to 150 μm for oocytes, with height maintaining an aspect ratio of 1.3). Polyacrylamide precursor solution including acrylamide/bis-acrylamide (7–12%T) and 3 mM BPMAC was degassed with sonication for 9 min. 0.08% APS and 0.08% TEMED were added to precursor solution and solution was pipetted between the SU-8 wafer (rendered hydrophobic with Gel Slick^TM^ solution) and a glass microscope slide functionalized with 3- (trimethoxylsilyl)propyl methacrylate (to ensure covalent grafting of PA gel to glass surface). After chemical polymerization (20 min), devices (glass with grafted PA gel layer) were lifted from wafer, rinsed with deionized water and stored dry until use.

### Mouse Embryo Isolation and Culture

Three-to-five-week-old female C57BL/6 J mice (000664, Jackson Laboratory) were superovulated by intraperitoneal (IP) injection of 5IU of Pregnant Mare Serum Gonadotropin (PMSG, Calbiochem, 367222, Millipore) and 46–48 hours later, IP injection of 5IU Human Chorion Gonadotropin (hCG, Calbiochem, 230734, Millipore). Superovulated females were housed at a 1:1 ratio with 3–8 month old C57BL/6 J males to generate 1-cell zygotes at 0.5 days post coitum. Using forceps under a stereomicroscope (Nikon SMZ-U or equivalent), the ampulla of oviduct was nicked, releasing fertilized zygotes and oocytes associated with surrounding cumulus cells into 50 μl M2 + BSA media (M2 media (MR-015-D, Millipore) supplemented with 4 mg/mL bovine serum albumin (BSA, A3311, Sigma)). Using a handheld pipette set to 50 μL, zygotes were dissociated from cumulus cells after the cumulus oocyte complexes were transferred into a 200 μl droplet of Hyaluronidase/M2 solution (MR-051-F, Millipore), incubated for up to 2 min, and passed through five washes in the M2+BSA media to remove cumulus cells.

From this point on, embryos were manipulated using a mouth-controlled assembly consisting of a capillary pulled from glass capillary tubes (P0674, Sigma) over an open flame attached to a 15-inch aspirator tube (A5177, Sigma). Embryos were passed through five washes of M2+BSA to remove cumulus cells. Embryos were then transferred to KSOM + BSA media (KCl-enriched simplex optimization medium with amino acid supplement, ZEKS-050, Zenith Biotech) that was equilibrated to final culturing conditions at least 3–4 hr prior to incubation. Embryos were cultured in 20 μl droplets of KSOM + BSA overlaid with mineral oil (ES-005-C, Millipore) in 35 × 10 mm culture dishes (627160, CellStar Greiner Bio-One) in a water-jacketed CO_2_ incubator (5% CO_2_, 37 °C and 95% humidity).

### Single-Embryo Quantitative Reverse Transcription Polymerase Chain Reaction (qRT-PCR)

All single embryo cDNA was prepared using a modified version of the Single Cell-to-Ct qRT-PCR kit (4458236, Life Technologies). Whole embryos were isolated at the desired developmental stage. Using a mouth pipette, embryos were then passed through three washes of PBS. With a hand-held pipette set to 1 μL, a single embryo was collected and transferred to one tube of an 8 well PCR strip. Presence of embryo was visually confirmed in each tube prior to cDNA synthesis. To account for the larger sample input, twice the protocol recommended volume of Lysis/DNAse (20 μL) was added to each embryo and allowed to incubate at room temperature (RT) for 15 minutes. Then, 2 μL of Stop Solution was added and incubated for 2 min. At this point, half of the reaction was stored in −80oC conditions as a technical replicate and the remaining sample (11 μL) continued through the original Single Cell-to-Ct protocol. All qRT-PCR reactions were performed using SSO Universal SYBR Green SuperMix, as per manufacturer instructions (1725275, BioRad). Primer sequences used were Rfx1 (5’AGT GAG GCT CCA CCA CTG GCC G, 5’TGG GCA GCC GCT TCT C), Dicer-1 (5’GGA TGC GAT GTG CTA TCT GGA, 5’GCA CTG CTC CGT GTG CAA) and Dicero (5’CTC TTT CCT TTG AAT GTA CAG CTA C, 5’CAG TAA GCA GCG CCC CTC). All qRt-PCR analyses were performed on the StepOnePlus Real Time PCR system (437660, Thermo).

### Single-Embryo and Single-Blastomere Microfluidic Immunoblotting

Once the desired developmental stage was reached, embryos were transferred to a ~50 μL droplet of acid Tyrode’s Solution (T1788, Sigma) and incubated at 37oC for up to 2 min to remove the *zona pellucida*. If dissociation into blastomeres was required, embryos were first incubated with a 1:1 solution of Accutase^®^ and 10X trypsin (15090046, Thermo) at 37oC (time varied with development stage, ranging from 5 min for two-cell embryos to up to 5 hr for blastocysts). Embryos were then mechanically disrupted by passing embryo through capillary repeatedly until dissociation. Single embryos or blastomeres were washed with PBS and deposited into microwells of the PA gel. Microwells were imaged by brightfield microscopy to collect data on size, morphology and ensure occupancies of one embryo or blastomere per microwell. Lysis conditions, including buffer composition, temperature and treatment time, were optimized for each developmental stage (Table S1). Electrophoresis was performed at 40 V/cm for varying times (from 20 to 60 s, depending on developmental stage and protein targets). Immobilization of proteins by photocapture was carried out by illumination with UV light source (100% power, 45 s, Lightningcure LC5, Hamamatsu). Gels were washed in 1X TBST for at least 1 hr before probing with antibodies. Primary antibodies were incubated at 1:10 dilution (40 μL/gel, in 2% BSA in 1X TBST), while fluorophore-conjugated secondary antibodies were incubated at 1:20 dilution (40 μL/gel, in 2% BSA in TBST). In order to strip bound antibodies and reprobe for new targets, gels were treated with 2% SDS, 0.8% β-mercaptoethanol and 62.5 mM Tris base at 55oC for three hours, washed in TBST (1 hr) twice and then reprobed.

### Antibodies

Antibodies employed for analysis of embryos include: rabbit anti-β-Tubulin (Abcam, ab6046, polyclonal, LOT: GR31927-5), mouse anti-Dicer-1 (Santa Cruz, sc-136981, A-2, LOT: I1817), mouse anti-CDX-2 (Abcam, ab157524, CDX2-88, LOT: GR300552-6), rabbit anti-SOX-2 (EMD Millipore, AB5603, polyclonal, LOT: NG1863962), goat anti-GAPDH (Sigma, SAB2500450, polyclonal, LOT: 6377C2), rabbit anti-GADD45a (Invitrogen, MA5-17014, D.81.E, LOT: R12274975). Donkey secondary antibodies AlexaFluor 647-conjugated anti-mouse (A31571), AlexaFluor 594-conjugated anti-mouse (A21203) and AlexaFluor 488-conjugated anti-rabbit (A21206) were purchased from ThermoFisher Scientific CA, USA.

### Image Processing, Signal Quantification and Statistical Analysis

The datasets generated during and analyzed during the current study are available from the corresponding author on reasonable request. Statistical tests were performed using GraphPad Prism 7.0b. Quantification of fluorescence signal of protein immunoblots employed in-house scripts written in MATLAB (R2017a, Mathworks)^33^. Briefly, Gaussian curves were fitted to protein band fluorescence intensity profiles using MATLAB’s Curve Fitting Toolbox. Gaussian fit parameters of protein peak location and σ were used to compute area-under-the-curve (AUC) by integrating the fluorescence intensity profile for the peak width defined as 4σ. Fiji (ImageJ) was used to false-color fluorescence micrographs and overlay channels to create composite images. ImageJ was used to compute cell volume^50^. Briefly, cell boundaries were traced using the freehand selection tool. For area traces with circularity of > 0.9, we assumed spherical morphology, computed cell diameter (ϕ) from traced area and cell volume was calculated from the computed cell diameter.

## Acknowledgements

The authors acknowledge members and alumni of the Herr Lab for helpful discussions. Partial infrastructure support was provided by the QB3 Biomolecular Nanofabrication Center. This research was performed under a National Institutes of Health Training Grant awarded to the UCB/USCF Graduate Program in Bioengineering (5T32GM008155-29 to E.R.C.), a California Institute for Regenerative Medicine Predoctoral Fellowship (E.R.C.), an Obra Social “la Caixa” Fellowship (E.R.C.), a University of California, Berkeley Siebel Scholarship (E.R.C.), a National Science Foundation CAREER Award (CBET-1056035 to A.E.H.), National Institutes of Health grants (1R01CA203018 to A.E.H.; R01GM114414, R01CA139067, 2R01CA139067, 1R21HD088885 and R21HD088885 to L.H.), a Howard Hughes Medical Institute (HHMI) Faculty Scholar Award (L.H.), a Bakar Fellow award at UC Berkeley (L.H.), a research scholar award from the American Cancer Society (L.H.), and an F32 Postdoctoral Fellowship from the National Institutes of Health (CA192636-03 to A.J.M.).

## Author contributions

E.R.C. and A.J.M. conceived experiments and executed the experiments. E.R.C. performed immunoblotting experiments, analyzed immunoblotting data and produced all figures and tables; A.J.M. collected, cultured and handled mouse embryos, performed qRT-PCR experiments and analyzed qRT-PCR data. All authors wrote the manuscript.

## References

1. Motosugi, N., Bauer, T., Polanski, Z., Solter, D. & Hiiragi, T. Polarity of the mouse embryo is established at blastocyst and is not prepatterned. Genes Dev. 19, 1081–1092 (2005).

2. Alarcon, V. B. & Marikawa, Y. Unbiased Contribution of the First Two Blastomeres to Mouse Blastocyst Development. Mol. Reprod. Dev. 72, 354–361 (2005).

3. Fujimori, T., Kurotaki, Y., Miyazaki, J. & Nabeshima, Y. Analysis of cell lineage in two- and four-cell mouse embryos. Development 130, 5113–5122 (2003).

4. Biase, F. H., Cao, X. & Zhong, S. Cell fate inclination within 2-cell and 4-cell mouse embryos revealed by single-cell RNA sequencing. Genome Res. 24, 1787–1796 (2014).

5. Xue, Z. et al. Genetic programs in human and mouse early embryos revealed by single-cell RNA sequencing. Nature 500, 593–597 (2013).

6. Casser, E. et al. Totipotency segregates between the sister blastomeres of two-cell stage mouse embryos. Sci. Rep. 7, 1–15 (2017).

7. Torres-Padilla, M. E., Parfitt, D. E., Kouzarides, T. & Zernicka-Goetz, M. Histone arginine methylation regulates pluripotency in the early mouse embryo. Nature 445, 214–218 (2007).

8. Goolam, M. et al. Heterogeneity in Oct4 and Sox2 Targets Biases Cell Fate in 4-Cell Mouse Embryos. Cell 165, 61–74 (2016).

9. White, M. D. et al. Long-Lived Binding of Sox2 to DNA Predicts Cell Fate in the Four-Cell Mouse Embryo. Cell 165, 75–87 (2016).

10. Plachta, N., Bollenbach, T., Pease, S., Fraser, S. E. & Pantazis, P. Oct4 kinetics predict cell lineage patterning in the early mammalian embryo. Nat. Cell Biol. 13, 117–123 (2011).

11. Shi, J. et al. Dynamic transcriptional symmetry-breaking in pre-implantation mammalian embryo development revealed by single-cell RNA-seq. Development 142, 3468–3477 (2015).

12. Bischoff, M., Parfitt, D.-E. & Zernicka-Goetz, M. Formation of the embryonic-abembryonic axis of the mouse blastocyst: relationships between orientation of early cleavage divisions and pattern of symmetric/asymmetric divisions. Development 135, 953–962 (2008).

13. Piotrowska-Nitsche, K. & Zernicka-Goetz, M. Spatial arrangement of individual 4-cell stage blastomeres and the order in which they are generated correlate with blastocyst pattern in the mouse embryo. Mech. Dev. 122, 487–500 (2005).

14. Zheng, Z., Li, H., Zhang, Q., Yang, L. & Qi, H. Unequal distribution of 16S mtrRNA at the 2-cell stage regulates cell lineage allocations in mouse embryos. Reproduction 151, 351–367 (2016).

15. Bordeaux, J. et al. Antibody validation. Biotechniques 48, 197–209 (2010).

16. Trenchevska, O., Nelson, R. W. & Nedelkov, D. Mass spectrometric immunoassays for discovery, screening and quantification of clinically relevant proteoforms. Bioanalysis 8, 1623–1633 (2016).

17. Schnell, U., Dijk, F., Sjollema, K. A. & Giepmans, B. N. G. Immunolabeling artifacts and the need for live-cell imaging. Nat. Methods 9, 152–158 (2012).

18. Teves, S. S. et al. A dynamic mode of mitotic bookmarking by transcription factors. Elife 5, 1–24 (2016).

19. Zhu, Y. et al. Nanodroplet processing platform for deep and quantitative proteome profiling of 10 – 100 mammalian cells. Nat. Commun. DOI: 10.1038/s41467-018-03367-w (2018).

20. Gao, Y. et al. Protein Expression Landscape of Mouse Embryos during Pre-implantation Development. Cell Rep. 21, 3957–3969 (2017).

21. Hughes, A. J. et al. Single-cell western blotting. Nat. Methods 11, 749–55 (2014).

22. Kang, C.-C. et al. Single cell–resolution western blotting. Nat. Protoc. 11, 1508–1530 (2016).

23. Kang, C. C., Lin, J. M. G., Xu, Z., Kumar, S. & Herr, A. E. Single-cell western blotting after whole-cell imaging to assess cancer chemotherapeutic response. Anal. Chem. 86, 10429–10436 (2014).

24. Kim, J. J., Sinkala, E. & Herr, A. E. High-selectivity cytology via lab-on-a-disc western blotting of individual cells. Lab Chip 17, 855–863 (2017).

25. Tsichlaki, E. & Fitzharris, G. Nucleus downscaling in mouse embryos is regulated by cooperative developmental and geometric programs. Sci. Rep. 6, 1–7 (2016).

26. Epifano, O., Liang, L. F., Familari, M., Moos, M. C. & Dean, J. Coordinate expression of the three zona pellucida genes during mouse oogenesis. Development 121, 1947–1956 (1995).

27. Martín-Coello, J., González, R., Crespo, C., Gomendio, M. & Roldan, E. R. S. Superovulation and in vitro oocyte maturation in three species of mice (Mus musculus, Mus spretus and Mus spicilegus). Theriogenology 70, 1004–1013 (2008).

28. Marangos, P. in Oogenesis: Methods and Protocols (ed. Nezis, I. P.) 209–215 (Springer New York, 2016).

29. Chen, S., Lee, B., Lee, A. Y. F., Modzelewski, A. J. & He, L. Highly efficient mouse genome editing by CRISPR ribonucleoprotein electroporation of zygotes. J. Biol. Chem. 291, 14457–14467 (2016).

30. Mozziconacci, J., Sandblad, L., Wachsmuth, M., Brunner, D. & Karsenti, E. Tubulin dimers oligomerize before their incorporation into microtubules. PLoS One 3, 1–8 (2008).

31. Strumpf, D. et al. Cdx2 is required for correct cell fate specification and differentiation of trophectoderm in the mouse blastocyst. Development 132, 2093–2102 (2005).

32. Zhang, S. Sox2, a key factor in the regulation of pluripotency and neural differentiation. World J. Stem Cells 6, 305 (2014).

33. Sinkala, E. et al. Profiling protein expression in circulating tumour cells using microfluidic western blotting. Nat. Commun. 8, (2017).

34. Padovan-Merhar, O. et al. Single Mammalian Cells Compensate for Differences in Cellular Volume and DNA Copy Number through Independent Global Transcriptional Mechanisms. Mol. Cell 58, 339–352 (2015).

35. Kempea, H., Schwabeb, A., Crémazya, F., Verschurea, P. J. & Bruggemanb, F. J. The volumes and transcript counts of single cells reveal concentration homeostasis and capture biological noise. Mol. Biol. Cell 26, 797–804 (2015).

36. Jeong, J.-K. et al. Evaluation of reference genes in mouse preimplantation embryos for gene expression studies using real-time quantitative RT-PCR (RT-qPCR). BMC Res. Notes 7, 675 (2014).

37. Revil, T., Gaffney, D., Dias, C., Majewski, J. & Jerome-Majewska, L. A. Alternative splicing is frequent during early embryonic development in mouse. BMC Genomics 11, 399 (2010).

38. Pan, Q., Shai, O., Lee, L. J., Frey, B. J. & Blencowe, B. J. Deep surveying of alternative splicing complexity in the human transcriptome by high-throughput sequencing. Nat. Genet. 40, 1413–1415 (2008).

39. Wang, E. T. et al. Alternative isoform regulation in human tissue transcriptomes. Nature 456, 470–476 (2008).

40. Flemr, M. et al. A retrotransposon-driven dicer isoform directs endogenous small interfering rna production in mouse oocytes. Cell 155, 807–816 (2013).

41. Morris, S. A., Guo, Y. & Zernicka-Goetz, M. Developmental Plasticity Is Bound by Pluripotency and the Fgf and Wnt Signaling Pathways. Cell Rep. 2, 756–765 (2012).

42. Huh, D. & Paulsson, J. Random partitioning of molecules at cell division. Proc. Natl. Acad. Sci. 108, 15004–15009 (2011).

43. Schultz, R. M. Regulation of zygotic gene activation in the mouse. BioEssays 15, 531–538 (1993).

44. Nothias, J. Y., Miranda, M. & DePamphilis, M. L. Uncoupling of transcription and translation during zygotic gene activation in the mouse. EMBO J. 15, 5715–25 (1996).

45. Darmanis, S. et al. Simultaneous Multiplexed Measurement of RNA and Proteins in Single Cells. Cell Rep. 14, 380–389 (2016).

46. Schwanhausser, B. et al. Corrigendum: Global quantification of mammalian gene expression control. Nature 495, 126–127 (2013).

47. Snider, N. T. & Omary, M. B. Post-translational modifications of intermediate filament proteins: Mechanisms and functions. Nat. Rev. Mol. Cell Biol. 15, 163–177 (2014).

48. Macaulay, I. C., Ponting, C. P. & Voet, T. Single-Cell Multiomics: Multiple Measurements from Single Cells. Trends Genet. 33, 155–168 (2017).

49. Zischewski, J., Fischer, R. & Bortesi, L. Detection of on-target and off-target mutations generated by CRISPR/Cas9 and other sequence-specific nucleases. Biotechnol. Adv. 35, 95–104 (2017).

50. Johnson, D. E., Ostrowski, P., Jaumouillé, V. & Grinstein, S. The position of lysosomes within the cell determines their luminal pH. J. Cell Biol. 212, 677–692 (2016).

